# MentaLiST – A fast MLST caller for large MLST schemes

**DOI:** 10.1101/172858

**Authors:** Pedro Feijao, Hua-Ting Yao, Dan Fornika, Jennifer Gardy, Will Hsiao, Cedric Chauve, Leonid Chindelevitch

## Abstract

MLST (multi-locus sequence typing) is a classic technique for genotyping bacteria, widely applied for pathogen outbreak surveillance. Traditionally, MLST is based on identifying sequence types from a small number of housekeeping genes. With the increasing availability of whole-genome sequencing (WGS) data, MLST methods have evolved toward larger typing schemes, based on a few hundred genes (core genome MLST, cgMLST) to a few thousand genes (whole genome MLST, wgMLST). Such large-scale MLST schemes have been shown to provide a finer resolution and are increasingly used in various contexts such as hospital outbreaks or foodborne pathogen outbreaks. This methodological shift raises new computational challenges, especially given the large size of the schemes involved. Very few available MLST callers are currently capable of dealing with large MLST schemes.

We introduce MentaLiST, a new MLST caller, based on a *k*-mer voting algorithm and written in the Julia language, specifically designed and implemented to handle large typing schemes. We test it on real and simulated data to show that MentaLiST is faster than any other available MLST caller while providing the same or better accuracy, and is capable of dealing with MLST scheme with up to thousands of genes while requiring limited computational resources. MentaLiST source code and easy installation instructions using a Conda package are available at https://github.com/WGS-TB/MentaLiST.

## 1 Background

Since it was introduced by Maiden *et al.* in 1998 [Ma98], Multi-Locus Sequence Typing (MLST) has become a fundamental technique for classifying bacterial isolates into strains. It has been applied in a large number of contexts, especially related to pathogen outbreak surveillance [PLACN17]. MLST works by associating to an isolate a sequence type defined by a specific allelic profile based on an established MLST scheme. There exist MLST schemes for many important pathogens [Pu]. Prior to the use of Whole-Genome Sequencing (WGS) data, MLST schemes were based on a small number of carefully selected housekeeping genes (usually fewer than 10), which made the approach both portable with respect to first-generation sequencing technologies (mostly Sanger sequencing) and able to accommodate pathogen evolutionary modes that might confound evolutionary analysis, especially lateral gene transfer [PLACN17].

Recently however, several case studies illustrated that the reliance on a small MLST scheme might not provide enough resolution to separate isolates into epidemiologically meaningful clusters. For example, Jolley *et al.* showed that traditional MLST schemes were not able to discriminate separate sublineages within a clonal complex of *Neisseria meningitidis* [Jo12]. This observation has come at a time when advances in sequencing technologies and protocols have had a major impact on public health, as it is now common to rapidly obtain WGS data from a pathogen outbreak, allowing for monitoring at an unprecedented level of resolution [GL14, Kw15, Ro16, Le14, Ly16, GG16, DF16, Ly16, Ro16, Le14, Pi15, GL14, MRF14]. In the specific case of MLST, this has led to the emergence of MLST schemes based on a larger set of genes, such as *core genome* MLST (cgMLST), that consider the set of core genes shared by a group of related strains (generally a few hundred genes), and even *whole genome* MLST (wgMLST) schemes that rely on a set of thousands of genes, covering most of the loci of the considered isolates [Ma13]. Since their introduction, cgMLST and wgMLST have proven to be valuable typing methods in many studies [Ko14, Be15, Ru15, Me16, Go17, Ko14, Ma16] and are expected to become standard approaches for pathogen surveillance [DdBH16].

These new developments in pathogen isolate genotyping motivate the development of MLST software able to accurately classify isolates into sequence types from large-scale WGS data that scale well in terms of the computational resources required, in order to handle large MLST schemes. Very few MLST tools that can meet these requirements currently exist. Indeed, most MLST typing programs have been developed with small schemes in mind, and are based on the availability of assembled genome sequences for the isolates being considered, or, if WGS data are provided, require an initial stage of contig assembly prior to the specific genotyping phase [Hu17, Br17, La12, PPP15, ml, Yo16, Pi15]. This approach suffers from the computational cost of assembling genomes, but more importantly for large MLST schemes, from the fact that reads corresponding to some loci in the scheme might not be assembled into contigs due to depth of coverage or other assembly issues. Other approaches have been developed recently that bypass the need to have assembled genomes or contigs and rely on directly mapping short reads onto the allele database for a given MLST scheme [In14, Te16]. However, the initial mapping phase is costly, especially for large schemes that can contain tens of thousands of alleles. Lastly, a few recent approaches have tried to avoid costly preprocessing of short read datasets by working on the principle of *k*-mer indexing, which has been shown to be helpful in handling large short read datasets in other bioinformatics contexts such as metagenomics [WS14]. Two tools that follow this approach currently exist: stringMLST [GJR16] and StrainSeeker [Ro17], although the latter assigns isolates to the nodes of a guide-tree which is required prior to the typing phase.

In this paper we introduce MentaLiST, a *k*-mer based MLST caller designed specifically for handling large MLST schemes. We test MentaLiST on several datasets, including a new cgMLST scheme for *Mycobacterium tuberculosis* composed of 553 essential genes, and compare its performance with that of two recent MLST callers, stringMLST, another *k*-mer based tool and ARIBA, a recent assembly-based tool. Our tests show that MentaLiST achieves comparable or better accuracy levels than both stringMLST and ARIBA while consistently using a low amount of memory and requiring much less computation time.

## 2 Methods

We propose a method for MLST calling that does not require pre-assembled genomes, working directly with the raw WGS data, and also avoids costly preprocessing steps, such as contig assembly or read mapping onto a reference. MentaLiST uses an algorithm that follows the general principle of *k*-mer counting, introduced in stringMLST [GJR16]. The first step in running MentaLiST is to create a *k*-mer hash map for a given a MLST scheme, mapping *k*-mers to the alleles in each locus where they are present; this hash map is then stored as a file. Next, MentaLiST can call the alleles for any given sample, by counting *k*-mers assigned to each allele, filtering out reads that do not reach a certain threshold. Each *k*-mer votes for all the alleles in which it is present in a locus, and the most voted allele from each locus is then chosen. Each step is detailed in the following subsections.

### 2.1 *k*-mer hash map

The first step is to build a *k*-mer database for a given MLST scheme and a given value of *k*. For each locus in the scheme, the *k*-mers of all alleles of the locus are computed, and a hash table linking each *k*-mer to the alleles where it is present is created. Each *k*-mer points to a list of loci, and for each locus there is also a list of all the alleles containing this *k*-mer. The same *k*-mer can be found in different loci, although the larger the value of *k*, the least likely this is to happen.

Since most alleles have very similar sequences, with small differences such as single nucleotide variation (SNV) or small indels, the large majority of *k*-mers are either present in most alleles, on in just a few (if they overlap a variable position). Therefore, instead of storing the list of all the alleles containing a *k*-mer, if this list contains more than half of the total number of alleles, its *complement* is stored instead. This requires us to also store a boolean variable indicating if a hit means that this particular *k*-mer is present (true) or absent (false) in the given allele list. As there are often loci with hundreds of alleles and MentaLiST is designed to handle MLST schemes with hundreds to thousands of genes, even with the additional space needed to store the extra boolean, this method decreases significantly the hash map memory footprint, while speeding up the allele calling phase since potentially a lot fewer alleles are involved in the voting step.

### 2.2 *k*-mer counting and voting

For a given sample, MentaLiST iterates through each sequenced read, *k*-merizing the read and checking on the *k*-mer hash map each *k*-mer hit. If the number of *k*-mer hits to the same locus is less than (*L – k*) · *α*, where *L* is the read length, *k* is the *k*-mer length, and *α* is a user-defined parameter, the read is discarded. In other words, *α* is the minimum proportion of the *k*-mers of a read that have to be found in the hash map for a read to be accepted as potentially mapping onto an allele of the MLST scheme. With this parameter, we aim to filter reads that do not belong to a particular locus of interest, but might contain some common *k*-mers, which in turn can introduce errors in the allele calling procedure. An optional read filtering step, as it was introduced in stringMLST, is also used in Mentalist.

We also implemented a faster way to discard reads that are not going to be from a gene present in the MLST scheme. This approach checks if the *k*-mer exactly in the middle of the read is present in the *k*-mer hash map. If not, the read is immediately discarded. This approach avoids processing every full read and matching it in the *k*-mer hash map, greatly speeding up the process specially on very high coverage datasets. We call this filter the “quick read filter”.

If a read passes the filtering step, then each of its *k*-mers gives one vote for all the alleles where this *k*-mer is present, as found through the hash map. The list might indicate the presence or absence of the given *k*-mer, as it was discussed above, so we give a weighted +1 or –1 vote, respectively, to all the alleles in the list. After all the reads have been processed, the allele with the largest vote count is selected for each locus. In the case of a tie, a random allele is selected among those with the most votes. MentaLiST outputs a log file with the number of votes for each allele on each locus and a list of tied alleles.

In the current version, MentaLiST expects that all genes in the MLST scheme are present in the given sample, since it is focuses on core genome MLST schemes.

### 2.3 Parameter calibration

The main parameters that can impact the behavior of MentaLiST are the *k*-mer length *k* and the read filtering threshold *α*. The default parameters were chosen through a calibration experiment with some simulated training datasets: MentaLiST was tested with thresholds varying from 0 to 0.8, and four *k* values: 17, 21, 25, and 31. As shown in Fig. 1, MentaLiST made fewer errors when *k* increased. No incorrect predictions were returned when the threshold was 0.3, no matter what the *k* value was. Hence, we assigned *α* = 0.3 to be the default threshold and *k* = 31 to be the default *k* value.

**Fig. 1:**
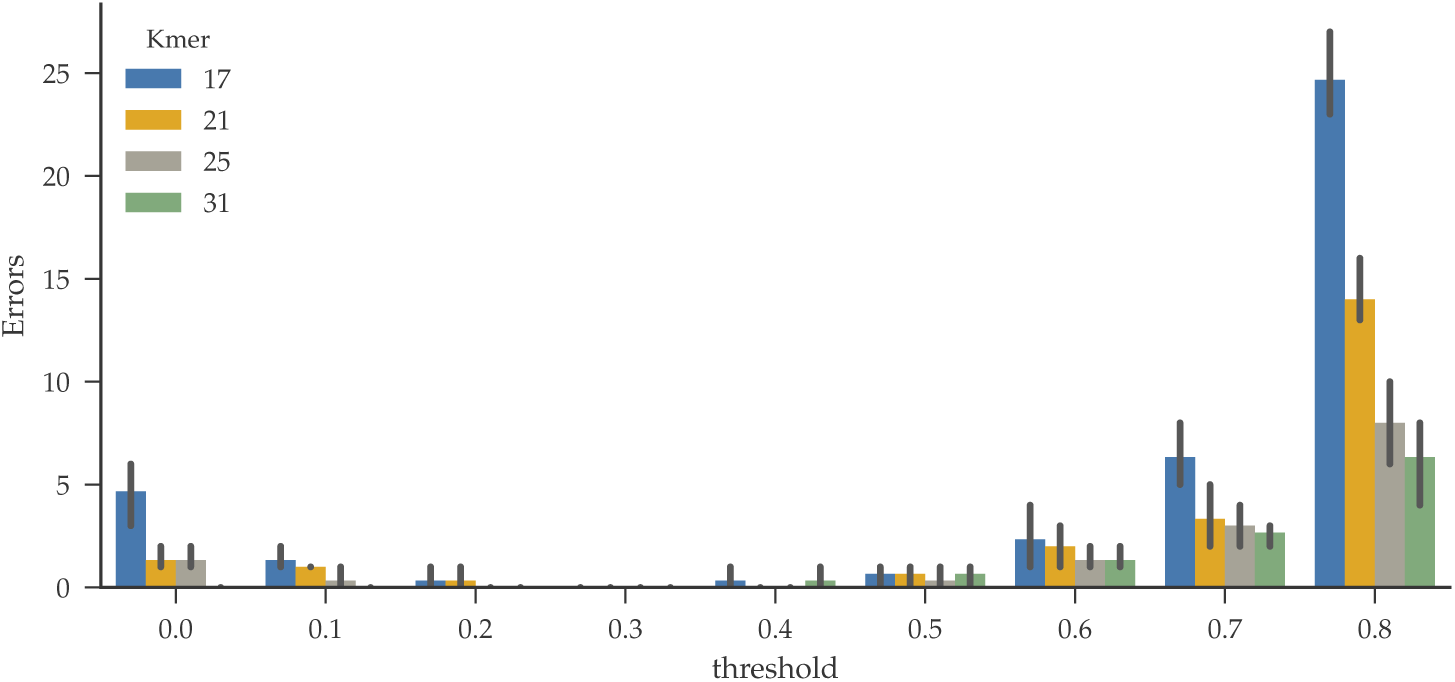
Average number of wrong calls on 3 simulated *M. tuberculosis* samples with different *k*-mer lengths and thresholds.

### 2.4 Experiment design

We evaluated the performance of MentaLiST on several real and simulated datasets, measuring both the calling accuracy, the running time and the computational resources required by MentaLiST. Following a recent review of MLST tools [Pa17], we compared MentaLiST with two top-performing MLST callers, ARIBA [Hu17], chosen for the accuracy of its calls, and stringMLST [Br17], that has been shown to be very fast. We ran two versions of MentaLiST, with the quick read filter enabled or disabled (referred to as MentaLiST Fast and MentaLiST, respectively).

The following datasets were used: The first dataset has 41 *Enterococcus faecium* samples, genotyped on a traditional MLST scheme based on seven housekeeping genes, taken from the ARIBA publication [Hu17].

The second dataset was based on *Mycobacterium tuberculosis* samples. We created an *essential core genome MLST scheme* (ecgMLST), which was based on selecting, from the full 2,891 genes in the cgMLST scheme from [cg], a subset of 553 essential genes, by intersecting the full cgMLST scheme gene set with the 615 essential *M. tuberculosis* genes described in [De17]. Since the loci in this ecgMLST scheme have gene annotations that match the *M. tuberculosis* H37Rv reference genome (NCBI accession number NC_000962.3), we constructed simulated genomes by substituting the reference sequence with a randomly chosen allele in each locus position.

We also considered two large *Salmonella* MLST schemes, the SISTR scheme with 330 genes [Yo16] and the Enterobase cgMLST scheme with 3002 genes [En]. In this case, since the annotations on both schema were not compatible with any *Salmonella* reference genome to the best of our knowledge, the simulated genomes were built by selecting a random allele for each locus and inserting random DNA between each pair of consecutive loci.

For all simulated genomes, WGS samples were created using the ART read simulator [Hu11], using the default parameters for an Illumina HiSeq 2500 machine. The samples were generated with read length and mean fragment size of 125/400 and 76/200 bp, respectively, with a 100x coverage in both cases. For the *M. Tuberculosis* dataset, we also created a mixed strain dataset, with samples containing reads from two strains with varying proportions, and a coverage test dataset, where samples have coverages from 10x to 100x, in increments of 10.

The tests were run on British Columbia Genome Sciences Centre’s computer cluster using a single thread for each application. We used the default value of the parameters for all three programs, except for the *k* value for stringMLST where it was fixed to *k* = 31, the same value as MentaLiST.

## 3 Results

### 3.1 *Enterococcus faecium* samples with a traditional MLST scheme

The first dataset we considered consists of 41 real *Enterococcus faecium* samples which have been used to evaluate ARIBA in its original publication [Hu17]. We used the *E. Faecium* MLST scheme downloaded from PubMLST [Pu] containing seven housekeeping genes. This experiment provides a comparison point for the use of MentaLiST on a classical, small-scale, MLST scheme.

As expected for a traditional MLST scheme, all tested methods made identical calls on all 41 samples. The main difference between the three programs was the runtime as shown in Fig. 5. ARIBA taking around 100 seconds per sample on average, was the fastest. On the other hand, stringMLST and MentaLiST both took around 300 seconds to make predictions, thus taking longer than but staying within the same order of magnitude as ARIBA. The runtime of MentaLiST Fast was shorter than stringMLST ‘s and MentaLiST ‘s on all 41 samples. The fast version spent less time than ARIBA on two samples, SRR98585 and SRR980586, characterized by a high coverage, 201x and 351x, respectively.

### 3.2 Essential cgMLST scheme for *Mycobacterium tuberculosis*

Our second experiment used the ecgMLST scheme for *Mycobacterium tuberculosis*, and focused on the impact of depth of coverage on the accuracy of type calls. We constructed three simulated *M. tuberculosis* datasets, and then for each one, we randomly selected reads to simulate a depth of coverage ranging from 10x to 100x in increments of 10x. All four programs, MentaLiST, MentaLiST Fast, stringMLST and ARIBA were tested in this setting, with the same parameterization as in the previous experiment.

MentaLiST, both in its standard version and in its fast version, was approximately one order of magnitude faster than stringMLST and three order of magnitude faster than ARIBA as shown in Fig. 5, at all coverage levels. MentaLiST Fast using less than 100 seconds, was the fastest application. ARIBA took more than 2 hours to assign samples to sequence types on this 553 genes scheme. As shown in Fig. 2, MentaLiST was the only application that accurately predicted all 553 genes on 10x coverage. stringMLST made one prediction error and provided no result for several genes. The calls for all programs were all correct when coverage was greater than 20x, except for stringMLST that returned one ambiguous call for only one gene, for some datasets.

**Fig. 2:**
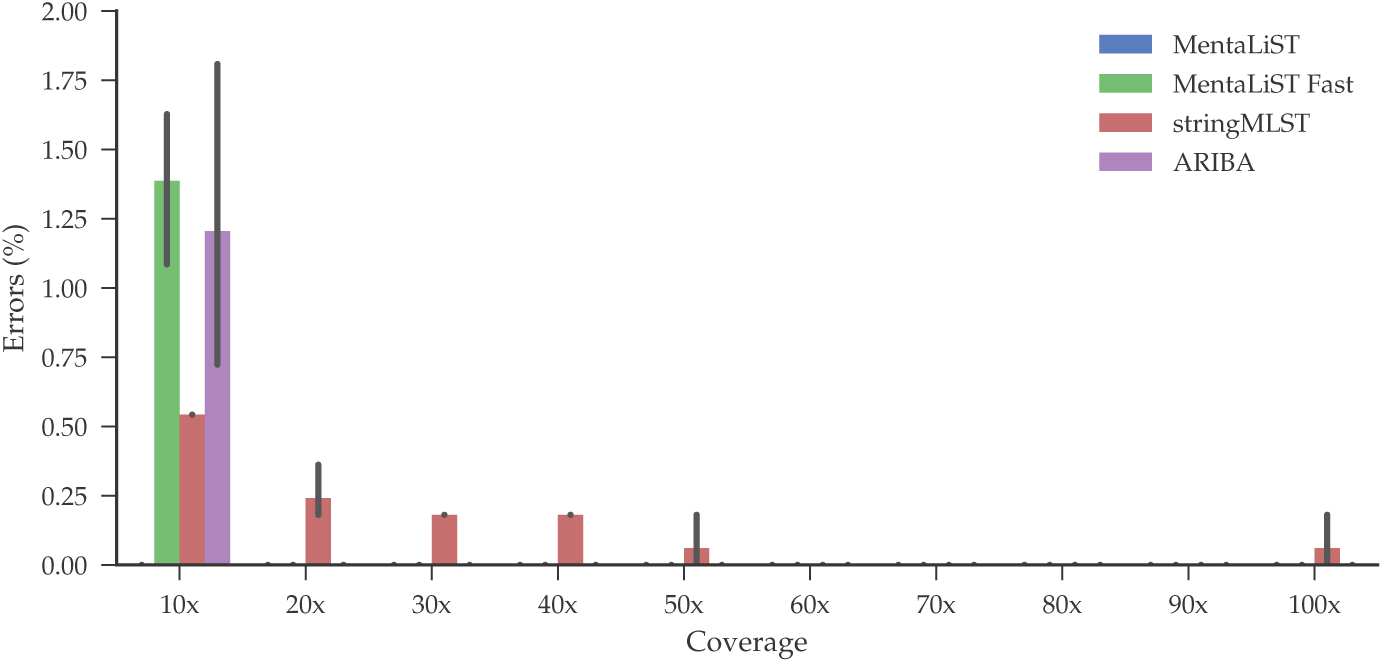
Average number of calling errors from three *M. tuberculosis* simulated samples, with varying depth of coverage and using the 553 genes ecgMLST scheme.

### 3.3 Mixed *M. tuberculosis* samples

Isolates from patients that suffer from an infection with more than one strain are not uncommon, with important pathological implications [Co12], and result in challenging WGS datasets, especially in terms of identifying the types present as well as their relative abundance [Ey13, Sa16]. To assess the performance of MentaLiST in that context, we generated simulated datasets composed of two strains at various levels of relative abundance. We generated three additional random strains, as described above, and mixed them with the previously described ones to generate three samples with two strains each, and relative abundance of the major strain, denoted by *p*_*major*_, ranging from 50% to 95%.

The run time results were similar to those of the previous experiment: MentaLiST was about 18 times faster (160 seconds per sample) than stringMLST (over 40 minutes per sample). MentaLiST Fast was still the fastest application with 80 seconds on average. However, MentaLiST Fast was less accurate than the two other methods, while MentaLiST and stringMLST performed similarly (see Fig. 3). They both returned correct calls when *p*_*major*_ was higher than 0.7. The only variation appeared when *p*_*major*_ was equal to 0.7. MentaLiST was free of error while stringMLST could not make a call for one gene. MentaLiST Fast and ARIBA ware more sensitive for the existence of the second sample. Correct calls were made when *p*_*major*_ is larger or equal to 0.75, showing a good robustness of all methods to mixed strain data.

**Fig. 3:**
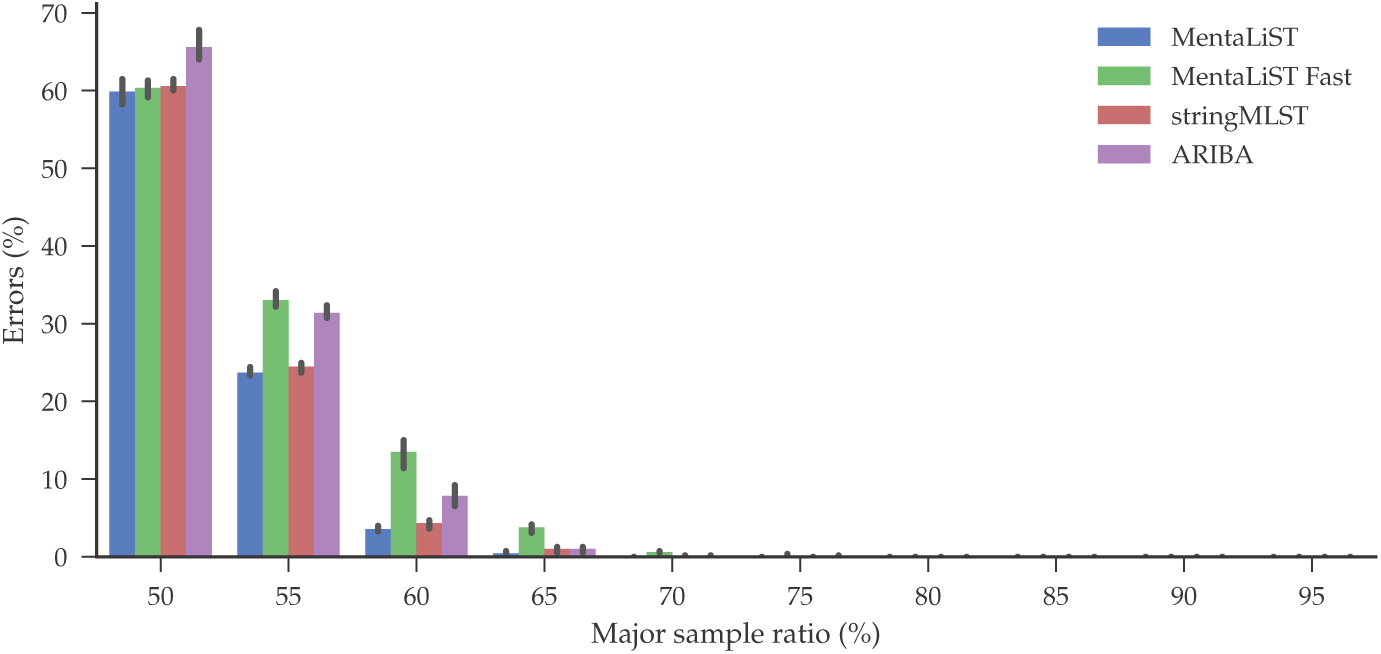
Average number of calling errors of three *M. tuberculosis* simulated datasets as a function of the proportion of the minor strain, using the 553-gene ecgMLST scheme.

### 3.4 Simulated *Salmonella* datasets

In the last test, we used simulated data of the Gram-negative bacterium *Salmonella* using two MLST schemes, the SISTR cgMLST scheme with 330 genes [Yo16] and the Enterobase cgMLST scheme with 3002 genes [En]. We generated ten genomes per scheme, and used two generated NGS datasets per genome with ART as previously detailed.

Since the Enterobase is a very large scheme, with 3002 genes, each with many alleles, stringMLST failed to build the alleles database and ARIBA failed to run because of memory errors. Hence, only the two MentaLiST versions were tested on this scheme.

Using the SISTR scheme, stringMLST and ARIBA took, respectively, ten and fifty minutes per sample to complete, roughly an order of magnitude slower than MentaLiST and MentaLiST Fast which took 85 seconds per sample on average (see Fig. 5). As shown in Fig. 4, ARIBA had the best performance, while making fewer errors in the 125bps datasets than the 76bps, whereas the opposite happened for the other callers. MentaLiST and MentaLiST Fast made one more error on average using reads of length 125bp than 76bp, with stringMLST having the worst performance. For the Enterobase scheme, regardless of the read length, about 30 of MentaLiST ‘s calls per sample were wrong, which equals a 1% error rate. Interestingly, the fast version made two fewer mistakes, and used only half the time that MentaLiST did.

**Fig. 4:**
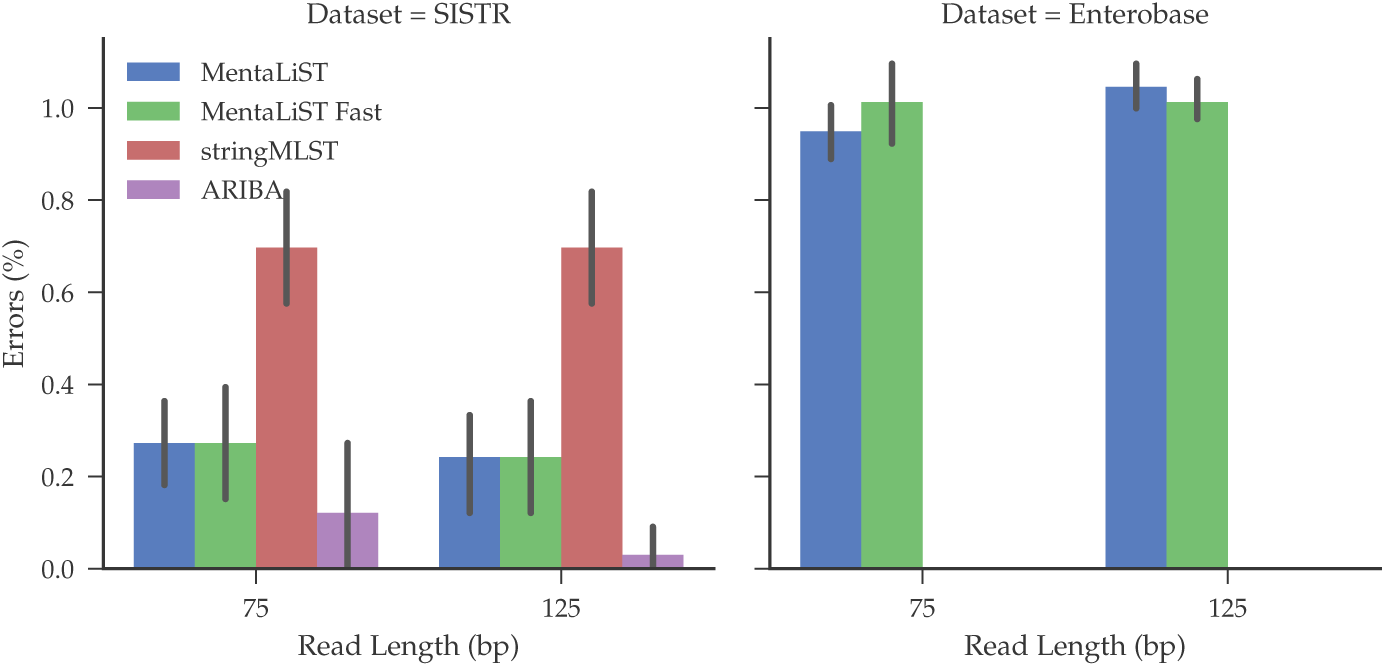
Average percentage of wrong calls on 10 simulated *Salmonella* samples with different read lengths for the SISTR (left) and Enterobase (right) cgMLST schema.

**Fig. 5:**
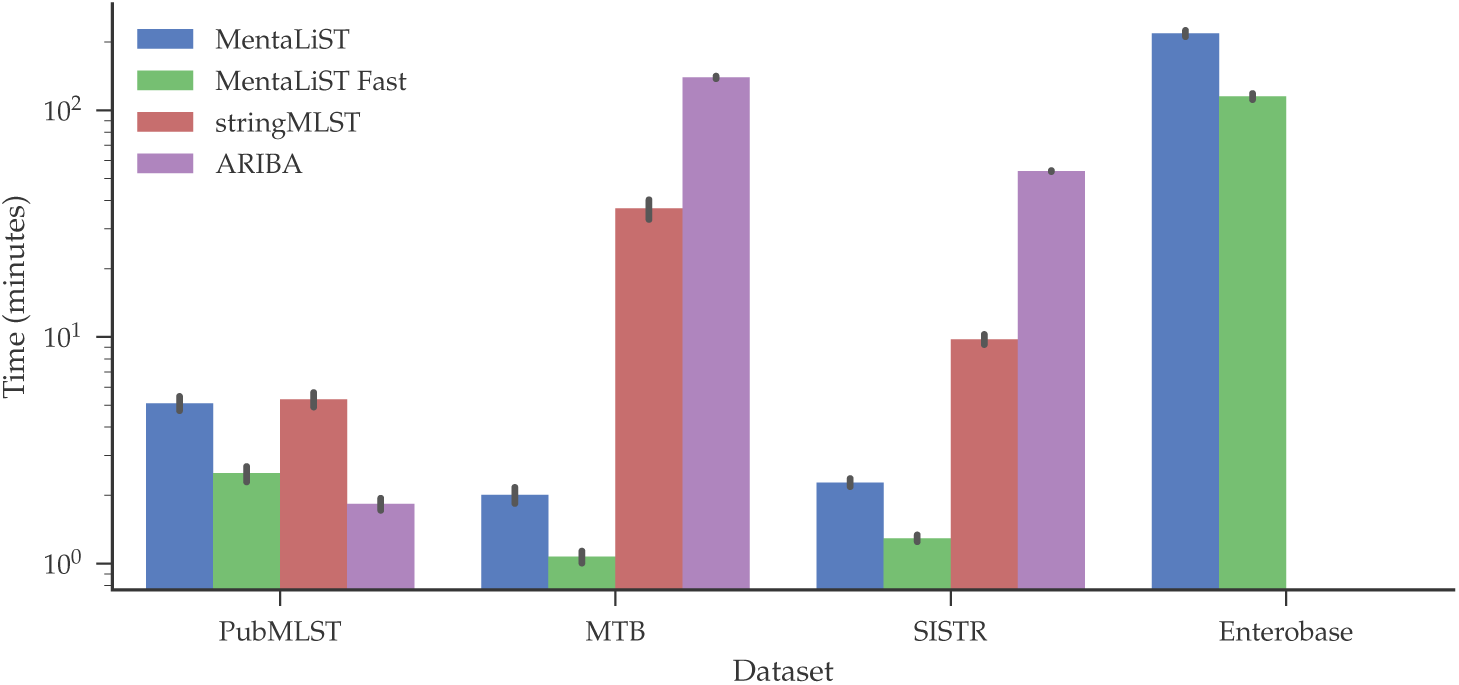
Running time for all programs on the different schemes.

**Fig. 6:**
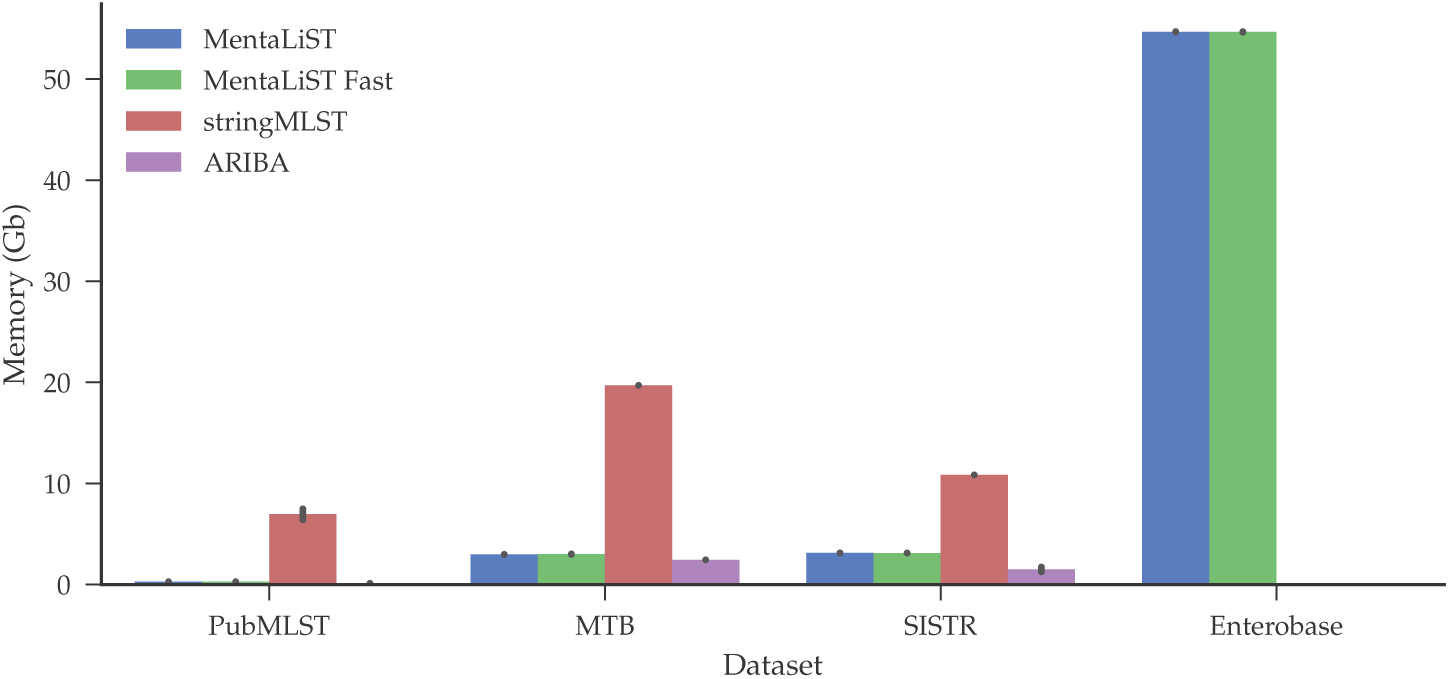
Peak memory usage for all programs on the different schemes.

## 4 Discussion

### 4.1 The importance of handling large MLST schemes

As demonstrated by our results, MentaLiST successfully handles large MLST schemes, including core-genome (cgMLST) schemes, up to a few thousand genes.

It has been previously recognized that these schemes are more accurate at identifying related strains in a surveillance context [Ko14, Me16, Go17] as they provide a higher resolution than the traditional schemes based on a handful of housekeeping genes.

While handling a scheme with thousands of genes and hundreds of alleles for each one is certainly taxing in terms of computational resource requirements, MentaLiST ‘s memory-efficient approach (using the simple, yet effective idea of storing the complement of a list of alleles containing a given *k*-mer when that list contains more than half of all the alleles for a given gene) greatly reduces both the time and the memory required for allele calling, without any adverse effect on the accuracy of the calls.

In addition, thanks to an efficient implementation in the Julia language [JBS17], MentaLiST significantly outperforms its closest competitors in terms of resource usage while providing the same or better accuracy, as we discuss in the following subsection.

### 4.2 Comparison with other MLST software

Since the majority of the existing publicly available software require either a mapping of the short reads onto a reference genome or their assembly into contigs as a preprocessing step, we did not compare our performance to theirs, as the amount of time and memory they would consume would likely make their practical application nearly-prohibitive on the kind of large-scale MLST schemes that we considered here. This left us with two tools in a comparable category-StrainSeeker [Ro17] and stringMLST [GJR16]. The former requires a guiding tree to be constructed on the alleles, and was thus excluded from the comparison. Instead, we added ARIBA [Hu17], which, although not necessarily designed to work on large MLST schemes, is nevertheless highly accurate.

Our results show that MentaLiST runs faster than stringMLST and uses less memory (due in part to the use of a better indexing scheme), while consistently providing the same or better call accuracy. Furthermore, this accuracy is always the same or better than that of ARIBA except on the 125-bp reads generated from the SISTR database [Yo16]. Therefore, our results enable us to confidently state that MentaLiST is at least comparable to the best-performing MLST callers in the class of the tools able to handle large MLST schemes with a reasonable amount of computational resources.

### 4.3 Future work

In future work, we would like to further investigate the possibility of using MentaLiST for determining the presence of a mixed infection, an important phenomenon in pathogens such as *M. tuberculosis*. Our experimental results in this setting (currently focused only on identifying the major strain) suggest that having a close, nearly-tied vote for best allele in several genes could be a signature of multiple strains being present in a sample at comparable frequencies.

In addition, if we were to allow ourselves a preprocessing step on the scheme such as the one required by StrainSeeker [Ro17], we could consider the use of a compression scheme using an approach inspired by pan-genomics such as Bloom Filter Tries [HWS16]. This could further reduce the memory requirements by adopting an even more efficient data structure for storing the *k*-mers found in the MLST scheme, and potentially speed up the voting process as well.

Reducing memory usage will allow the application of MentaLiST to whole genome MLST schemes. However, a new issue must also be taken into consideration, namely the detection of missing genes, present in the MLST scheme but not in the sample. We hypothesize that by comparing the number of expected matching *k*-mers for a gene present in the sample to the actual number of matches, it is possible to detect missing genes in cases where a significant difference is observed.

Lastly, an intriguing application of MentaLiST would be to identify and flag novel alleles that are not present in the scheme provided for a given gene. Currently, MentaLiST considers only those *k*-mers that are present in at least one allele in the existing scheme; however, if it is found that a sufficient number of reads fall slightly below the threshold they need to meet to participate in the voting process, this could indicate that a novel allele is present that might differ from an existing one in a small number of loci.

In conclusion, as the cost of WGS experiments continues to decrease while the number of sequenced organisms grows, tools such as MentaLiST will play an increasingly key role in relating bacterial sequences to one another, investigating their population-level diversity, and, most importantly, helping track the progression of infectious disease outbreaks.

